# Geographic-specific Diversification of Vaginal Microbiota in China

**DOI:** 10.1101/2024.11.08.622683

**Authors:** Hua Yang, Yina Liu, Zimo Liu, Chuanya Zhou, Lingjie Hu, Jintao He, Lingling Lv, Zhemin Zhou, Lan Zhu

## Abstract

The vaginal microbiota is pivotal to women’s reproductive health, with significant geographic and ethnic heterogeneity suggesting evolutionary adaptations to environmental, climatic, and lifestyle factors. In this multi-centric study, we analyzed vaginal microbiome samples from women across five provinces in China, each characterized by distinct climatic, ethnic, and lifestyle traits. Our results indicate that the vaginal microbiota among Chinese women is characterized by low alpha diversity, with *Lactobacillus crispatus* being the dominant species in Community State Type (CST) I, especially in the northern regions. While most participants exhibited stable CSTs throughout their menstrual cycles, fluctuations were observed in some individuals. We also detected low-abundance bacterial species such as *Fannyhesea vaginae* and *Sneathia vaginalis*. Comparative genomic analysis of high-quality metagenome-assembled genomes of *L. crispatus* revealed minimal single nucleotide polymorphism (SNP) variations within individuals but substantial diversity between individuals, delineating a distinct separation between vaginal-associated populations (VAP) and gut-associated populations (GAP). These findings highlight the intricate dynamics of the vaginal microbiome, which are markedly influenced by geographic and individual determinants. This study offers an in-depth phylogenetic and genomic analysis of the vaginal microbiota among Chinese women, underscoring the imperative for further research into the impact of regional and individual variations on microbial community composition. Such knowledge could facilitate the development of personalized vaginal health management strategies that accommodate the unique microbiome profiles specific to different geographical regions.

**Importance:** This study provides a groundbreaking analysis of the vaginal microbiota among Chinese women, offering insights into the population structure and evolutionary dynamics of this critical component of women’s reproductive health. The research reveals that *Lactobacillus crispatus*, a key species in vaginal health, has a recent evolutionary emergence and exhibits significant geographic variation in prevalence, being more common in northern China. This discovery, along with the identification of distinct vaginal-associated and gut-associated populations, underscores the importance of understanding the vaginal microbiota’s complexity. The findings have profound implications for developing personalized strategies to manage vaginal health, tailored to regional microbiome characteristics. This work not only advances our understanding of the vaginal microbiota’s role in health and disease but also sets the stage for future research into the factors that shape microbial community structures across diverse populations.

## Introduction

The vaginal microbiota plays a fundamental role in maintaining women’s reproductive health (1–3). This complex ecosystem consists of diverse microbial communities whose composition and structure are influenced by the host’s health, lifestyle, geography, and ethnicity (4–8). While the global microbiota has been categorized into five distinct community state types (CSTs) (9), dominated by species such as *Lactobacillus crispatus* (CST-I), *Lactobacillus iners* (CST-III), and *Gardnerella vaginalis* (CST-IV), the specific factors driving the evolution and geographical distribution of these microbial communities remain poorly understood.

Among the CSTs, *Lactobacillus crispatus* (CST I) is typically associated with optimal vaginal health, as its production of lactic acid and exopolysaccharides helps maintain a low vaginal pH, creating an inhospitable environment for pathogens (10–12). In contrast, CST IV, dominated by *Gardnerella vaginalis*, is often linked to dysbiosis and conditions like bacterial vaginosis (13). While some strains of *Gardnerella* coexist with the host without causing symptoms, others are pathogenic. The relationship between *L. iners* and vaginal health is more nuanced, as it often represents a transitional state between health and dysbiosis (14, 15).

Several studies have noted variations in vaginal microbiota composition based on ethnicity and geographic origin (6, 16, 17). For example, *Gardnerella* and *L. iners* are more prevalent in the vaginal microbiota of African American women, while *L. crispatus* is dominant in women of European ancestry (18). This geographic variation may reflect evolutionary processes that began during the migration of modern humans from Africa, with subsequent adaptations to local environments (19). Despite these insights, a comprehensive phylogenetic analysis of the dominant vaginal species in diverse populations, including China, is still lacking.

China is a vast country with diverse climates, ethnicities, and lifestyles, all of which could shape the vaginal microbiota (17, 18, 20). However, there is limited information on how these factors influence the composition and evolutionary history of vaginal bacteria across different regions of China. To address this gap, we conducted a multi-center study analyzing vaginal microbiome samples from five geographically distinct provinces. By reconstructing the genomes of the dominant bacterial species and comparing them with global data, we aimed to understand the population structure and evolutionary history of the vaginal microbiota in Chinese women. Using artificial intelligence (AI) models, we also identified key accessory genes and core genome mutations that may play pivotal roles in the microbial evolution within this population.

This study offers new insights into the geographic-specific evolution of the vaginal microbiota in Chinese women, highlighting the importance of both environmental and evolutionary factors in shaping microbial communities. Our findings provide a foundation for future research into the role of the vaginal microbiota in women’s health and may guide the development of region-specific treatments for vaginal dysbiosis.

## Results

### Temporal-spatial variation of vaginal microbiota in Chinese Population

A total of 30 healthy individuals across five provinces (Fig. 1A) were enrolled, and four vaginal mucosal samples were collected for each individual, during the menses, the follicular phase, pre-ovulation, post-ovulation, and luteal phases over one menstrual cycle. The majority of the samples were in low alpha-diversity, with an average Simpson diversity indices (SDIs) of 0.15 (0-0.82), indicating the presence of predominant species in most samples. The UMAP analysis separated all samples into four distinct clusters, each predominated by one species and associated with one described CST. CST-I (*L. crispatus*) was the most prevalent, accounting for 55/120 (45.8%) samples from 15 individuals, followed by CST-III (*L. iners*) with 42 (35%) samples from 12 individuals, and CST-IV B (*Gardnerella vaginalis*) with 19 (15.8%) samples from five individuals. Finally, CST IV C3, predominated by *Bifidobacterium breve*, was found only in one individual from Henan and remained stable throughout the menstrual cycle (Fig. 1B). Notably, all eight samples with an SDI of >0.55 were associated with an increase of *G. vaginalis* in vaginal microbiota, resulting in a significantly elevated SDI in CST IV B-associated samples (Fig. 1C).

**Figure 1.**
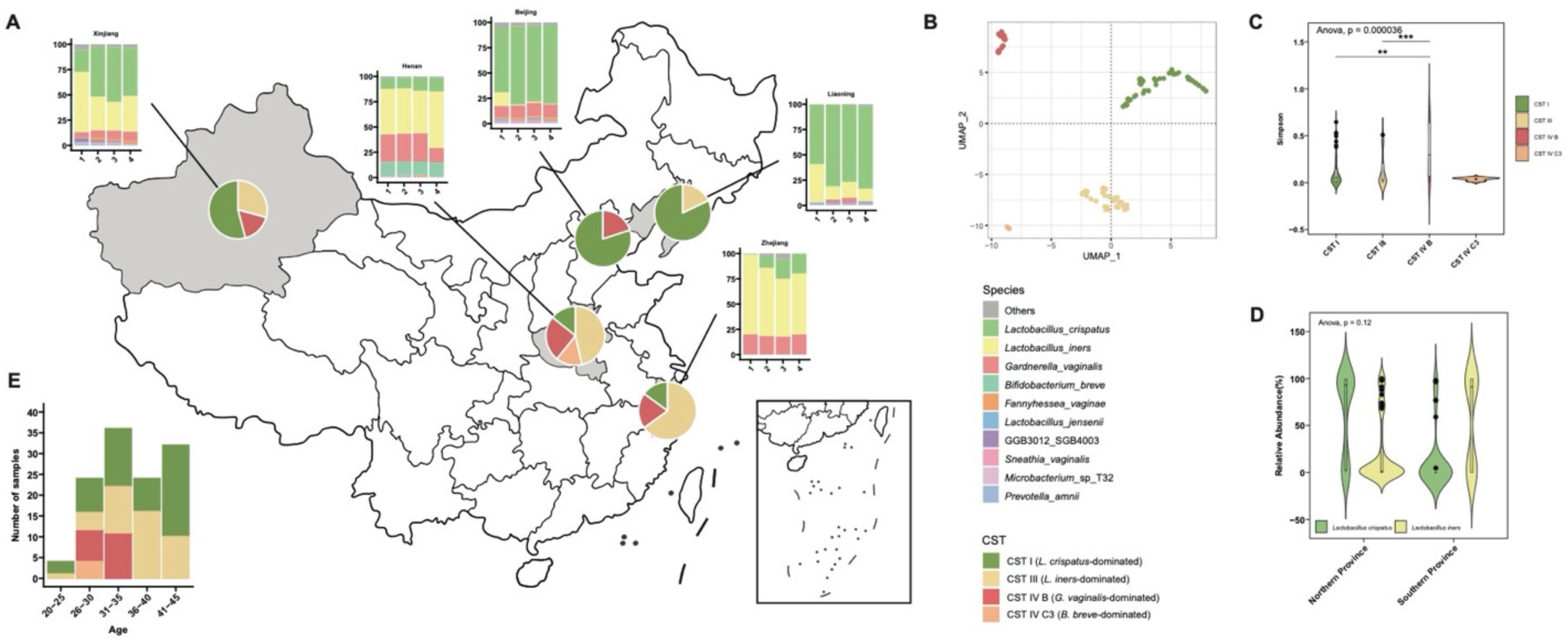
Community State Types (CST) and Species Composition of Chinese population A Geographical Distribution of CST and Species Composition(The bottom left shows the age distribution.The pie charts represent the percentage of different CST in various regions. The bar charts display the top 10 species in relative abundance from four different menstrual phases (follicular phase, pre-ovulation, post-ovulation, luteal phase) across different regions. B Group the CST types of the vaginal microbiome in the Chinese population using UMAP. C Simpson diversity values for different CSTs. D. Relative abundance of *Lactobacillus crispatus* and *Lactobacillus iners* in northern and southern provinces of China. E Age distribution of the samples.

Potential associations were identified between the CSTs and both the geography and the ages. The majority (60-80%) of the samples in Beijing, Liaoning, and Xinjiang were assigned to CST-I, significantly greater than the <20% CST-I samples in Henan and Zhejiang (p<0.0001), indicating a latitudinal-dependent distribution of *L. crispatus* (Fig. 1D). In contrast, CST-III were enriched in samples from southern provinces compared to those from Beijing and Liaoning. Notably, while the CST-IV types were of no geographic specificity and found in almost all regions except Liaoning, they were only detected in individuals between the ages of 26-35 (Fig. 1E).

The principal coordinates analysis (PCoA) revealed high similarities among samples from the same individuals, with an intra-person Bray-Curtiss distance of 0.16 versus the significantly greater distance (0.67) between different individuals (Fig. 2A). The majority of individuals remained in the same CST type throughout the menstrual cycle, except for three that converted from CST-III to CST-I before or post ovulation phase, and one that converted from CST-IV to CST-III before luteal phase (Fig. 2B). As a result, a gradual increase in *L. crispatus* or decrease in *G. vaginalis* were observed in all regions, towards overall healthier microbiomes over the menstrual cycle.

**Figure 2.**
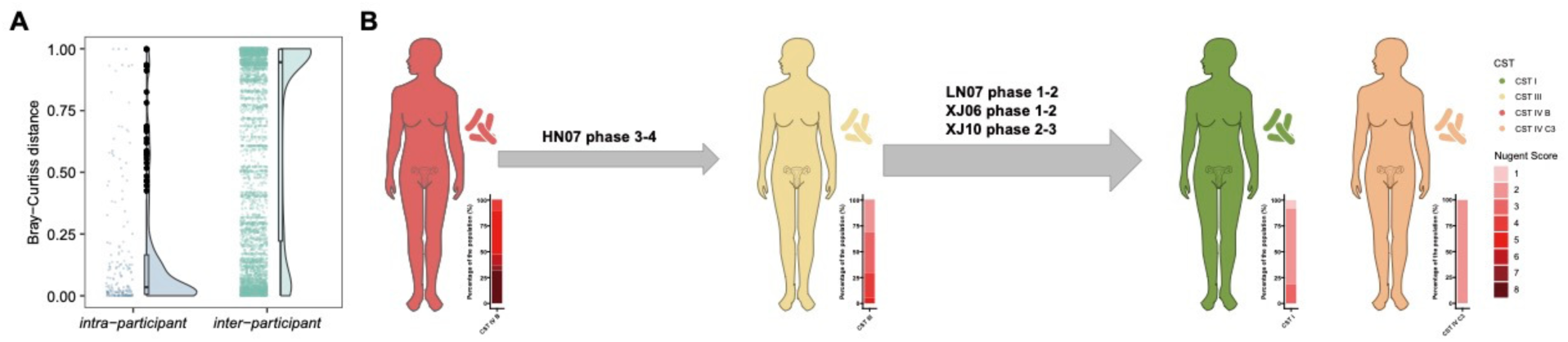
Changes in CST and Microbial Similarities Across the Menstrual Cycle A Principal coordinates analysis (PCoA) illustrating the Bray-Curtis distances between vaginal microbiota samples. Samples from the same individual exhibited high similarity, with an intra-person Bray-Curtis distance of 0.16, compared to a significantly greater inter-individual distance of 0.67. B Transition of individuals between CST throughout the menstrual cycle. The majority of individuals maintained the same CST type; however, three individuals converted from CST-III to CST-I before or during the follicular phase, and one individual transitioned from CST-IV to CST-III before the ovulation phase. Changes in *L. crispatus* levels and reductions in *G. vaginalis* were observed, indicating a trend towards healthier microbiomes over the menstrual cycle.

We also detected many other bacteria in the vaginal samples, albeit in low frequencies (Figure S 1). Over 1% of *Fannyhesea vaginae* and *Sneathia vaginalis* were found in ten CST-IV B samples from three individuals, while *Lactobacillus jensenii* and *Microbacterium* sp. were found in over half of the CST-I or CST-III samples, indicating inter-species collaborations of vaginal microbiota. Notably, frequent invasion of microbes from other sources was detected in samples of CST-III B, including a high level of *Prevotella corporis*, a typical oral bacteria, in XJ-04, 0.007% reads specific to *Bifidobacterium bifidum,* gut bacteria, and *Candida albicans*, a fungi, in BJ-03 and ZJ-01, respectively (Table S1).

### Geographic Distribution Characteristics of *Lactobacillus crispatus*

High-quality *L. crispatus* metagenome-assembled genomes (MAGs), with >5 read depths, were reconstructed from 67 samples in 17 individuals using Enterobase ToolKit (EToKi) (21). Notably, 59 of the MAGs were from the Northern provinces (Fig. 1A). The *L. crispatus* from the samples of the same individuals differed from each other by only 0-24 SNPs in the core genome. Further investigation revealed that almost all of these intra-individual single nucleotide polymorphisms (SNPs) were non-synonymous, resulting in amino acid variations in core genes (Table S2). The involved genes were *pepD* and *ypdF*. Intriguingly, the *L. crispatus* from different individuals also exhibited extremely high similarities in their core genomes. A total of 69 SNPs were identified among Chinese individuals, resulting in inter-personal diversities of only 0-44 SNPs in the core genome.

We integrated the reconstructed genomes with 316 publically available genomes to estimate a maximum likelihood phylogeny of *L*. crispatus (Fig. 3A), which was broadly separated into two of vagina- or gut-associated populations (VAP and GAP), following their different niche specificities. *L. crispatus* was likely lived in the guts, and later transferred into the vagina to form the VAP. This finding was further evidenced by the greater genetic diversity among strains in the GAP (w=0.02) than those in the VAP (0.007) (Fig. 3B). The bacteria might have experienced a host transfer in the progress, as the GAP was found in multiple mammals and avians whereas the VAP was exclusively found in humans.

**Figure 3.**
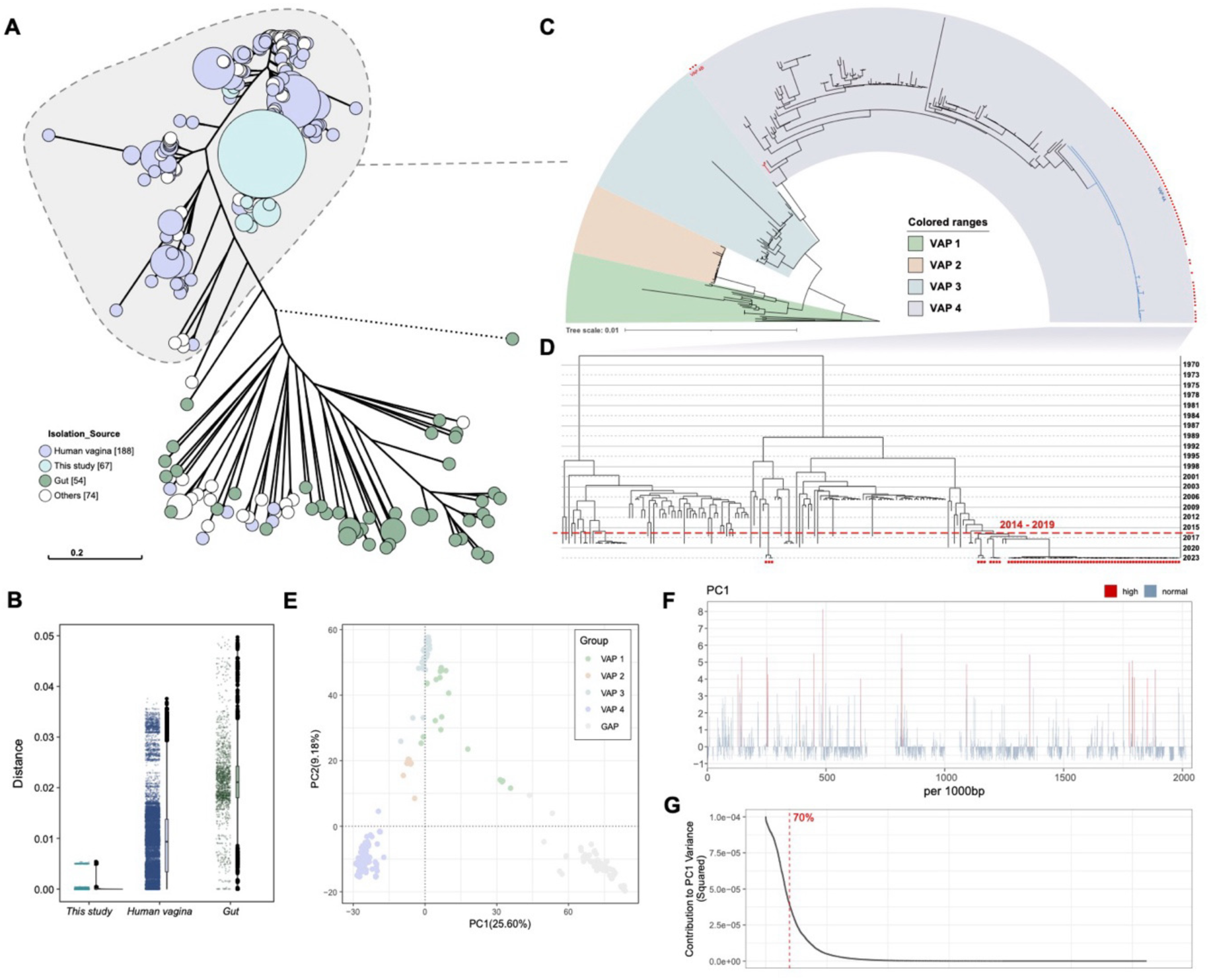
Analysis of the core genome of *Lactobacillus crispatus* A Maximum Likelihood Phylogenetic Tree of *Lactobacillus crispatus* (Includes *Lactobacillus crispatus* genomes obtained from this study and the NCBI public database). B Distribution of Distances Among *Lactobacillus crispatus* Strains (according to Figure A). C The vaginal associated population (VAP) phylogenetic tree constructed using hierBAPs, with different color ranges representing the four clusters annotated by hierBAPs. D The Bayesian phylogenetic tree constructed with the best model in BEAST2, with the red dashed line indicating the introduction time of the Chinese strains. E Principle Component Analysis (PCA) Constructing a Two-Dimensional Scatter Plot of the Core Genome of *Lactobacillus crispatus* Based on single nucleotide polymorphisms (SNPs): F Contribution of SNP Sites in *the Lactobacillus crispatus* Genome to PC1 (the horizontal axis represents every 1000 bp of the genome, using the sigma test. sites greater than 4σ are marked in red as significant). G The loading factors of each SNP site.

We separated the VAP into four clusters using hierBAPS, designated as VAP1 – VAP4, respectively(Fig. 3C). All Chinese strains fell into two clades (4A and 4B) within the VAP-4 cluster. VAP-4A consisted of samples from 64 individuals across 5 provinces, exhibiting no observable regional or individual specificity. Furthermore, two and one strains from South Korea and Italy also presented in the same cluster, demostrating its global presence. Additionally, all three samples in VAP-4B were from one individual in Xinjiang. We identified no more than three SNP differences for *L. crispatus* from the same clade, plus an inter-clade diversity of ∼0.004 (Fig. 3B), substantially lower than the average genetic diversities in VAP or GAP strains. To characterize the recent evolution of *L. crispatus* in China, we inferred the population dynamics of cluster VAP-4 using BEAST2(Fig. 3D). A total of 4 alternative models were compared using the nested sampling tests, and the model with optimal relaxed substitution rates and Bayesian skyline coalescents was selected with its greatest Bayes factor (Table S3). The analyses revealed a very recent origin of the VAP-4 cluster, dated in 1981 [confidence interval (CI) 95% 1976-1984]. This cluster experienced rapid population expansion in the past four decades, and the VAP-4A arrived in China in 2016 (VAP-4A; CI 95% 2014-2019). This clade was subsequently transmitted across China in 10-12 years and widely detected in all five provinces in this study. Furthermore, VAP-4B, only found in one individual, diverged from its closest neighbor from the US in 2006 (VAP-4B; CI 95% 2006-2006), possibly reflecting an occasional international transmission.

### Selective stress in the core-genome of *Lactobacillus crispatus*

We proposed an unsupervised approach to evaluate the impact of natural selection on the evolution of *L. crispatus*. To this end, we employed PCA analysis to embed all *L. crispatus* genomes in a 2-D space, where three clusters were visually identified, corresponding to GAP, VAP4, and VAP1-3, respectively (Fig. 3E). Furthermore, we measured the loading factors of each SNP site and found that the top 5.8% most contributing SNPs accounted for >70% of differences in PC1 (Fig. 3F). These SNPs had an average loading factor of 7.2e-5, 38X greater than that of the remaining SNPs (1.9e-6). We plotted all SNPs along the chromosome and identified 18 genes with greater average loading factors than other regions (sigma >= 4) using a sliding window of 1Kb (Figure 3G, Table S4). The variations in these regions significantly contributed to the separation of the two clusters. The genes under diversifying selection between different niches included *lea* (E6A57_01235), an *LEA* family epithelial adhesin that was known for adhesive of the bacteria on epithelial cells. Many other genes, including *gtfA*, encoding glucosyltransferase A, *murG*, encoding genes for peptidoglycan layer formation, *asnB*, encoding asparagine synthase, and E6A57_09405 for the alpha-glucosidase, were all associated with biofilm formation, which is essential for survival in vaginal epithelial cells.

### Analysis of the Pan-Genome of *Lactobacillus crispatus*

The pan-genome analysis revealed a substantially different evolutionary scenario in *L. crispatus*. Despite their 50% lower core genomic variations, the VAPs exhibited substantially greater variations than GAPs in the accessory genome, changing 22% of their accessory genes than 20% in GAPs (Fig. 4B). The Chinese samples, exhibiting almost no variations in their core genome, varied 8% in the accessory genome. They also formed a much looser cluster in the tree based on gene presence/absence (Fig. 4A). Furthermore, four samples fell outside of the Chinese cluster, indicating complicated gene acquisitions or losses. Detailed analysis indicated that these massive accessory genome variations were associated with the repetitive acquisition of mobile genetic elements, especially prophages and plasmids, possibly reflecting rapid adaptive radiation after its adaptation to the new niche.

**Figure 4.**
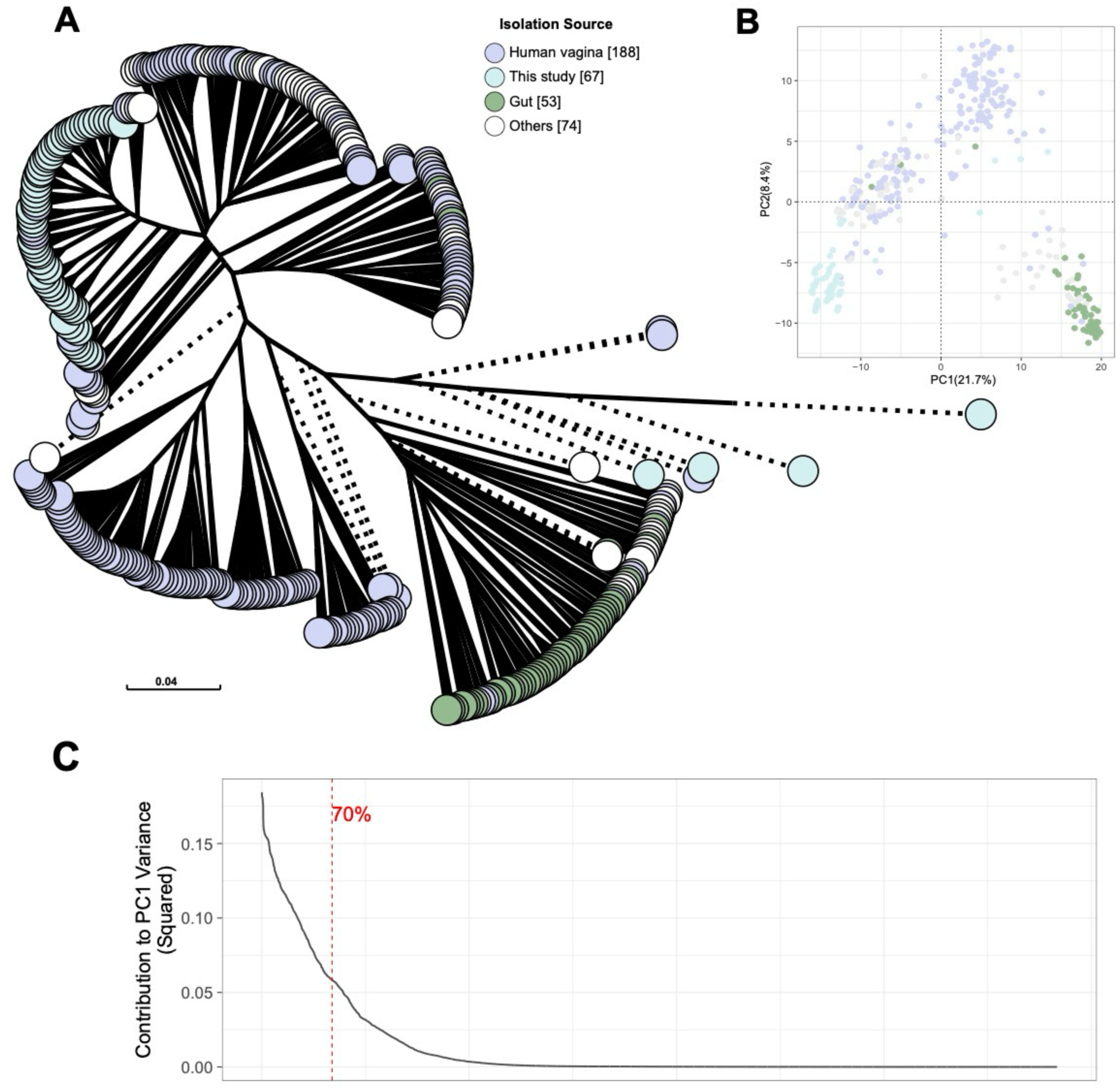
Pan-Genomic Analysis of *Lactobacillus crispatus* A Pangenome Tree of *Lactobacillus crispatus* (calculate the Bray-Curtis dissimilarity index between samples based on the pangenome gene presence/absence matrix and perform sample clustering analysis using the UPGMA method). B. PCA analysis constructing a two-dimensional scatter plot of the *Lactobacillus crispatus* pangenome based on the presence/absence of pangenome genes. C. PCA of *Lactobacillus crispatus* accessory genomes shows significant differences between VAPs and gut-associated populations (GAPs). Accessory genes (8.8%, 848/9586) contribute to 70% of the variation in PC1, indicating greater genomic differentiation in the pan-genome.

PCA analysis also revealed the substantial accessory gene differences between VAPs and GAPs. Different from the core genome SNPs, a much larger proportion of accessory genes (8.8%, 848/9586) contributed to 70% of the variations in the first PC, demonstrating greater inter-population variations in the pan-genome (Fig. 4C, Table S5).

Detailed analyses of the varied genes revealed an enrichment of nutrient-utilising genes in VAP, including *nanA* and *nanE*, which were associated with the acquisition of sialic acid in the cervical-vaginal fluid, and *agaA*, *sglT*, and *bglK*, involved in polysaccharide metabolism. Meanwhile, we also observed depletions of *glpF* and *dhaMLK* genes, responsible for the update of glycerol in the gut. Furthermore, VAP also lost *axeA* and *eutT*, responsible for the utilisation of *hemicellulose* and *cobalamin*, both enriched in the gut.

### Evolutionary Analysis of *Lactobacillus iners*

We also applied the same phylogenetic analyses on both *L. inners* and *Gardnerella vaginalis*, which exhibited distinct phylogenetic dynamics from *L. crispatus* (Fig. 5A). The *L. inners* genomes were genetically about 4X more diverse than *L. crispatus* (0.09 vs 0.02) globally, and the Chinese strains exhibited only slightly reduced core genomic variations than the international ones (0.043 vs 0.09) (Fig. 5B). PCA analysis also suggested a lack of genetic separation in the *L. inners* strains, although the Chinese strains were enriched at the bottom of the cluster (Fig. 5C). Phylogenetic analysis indicated that the Chinese *L. inners* strains from both this study and other, public projects fell into a loose cluster in the tree, together with many strains from the US and South Africa, indicating frequent historical, international transmissions. Notably, we observed geographic specificity in China, with the majority of samples from Southern regions located near the root of the cluster, whereas the northern regions were further away.

**Figure 5.**
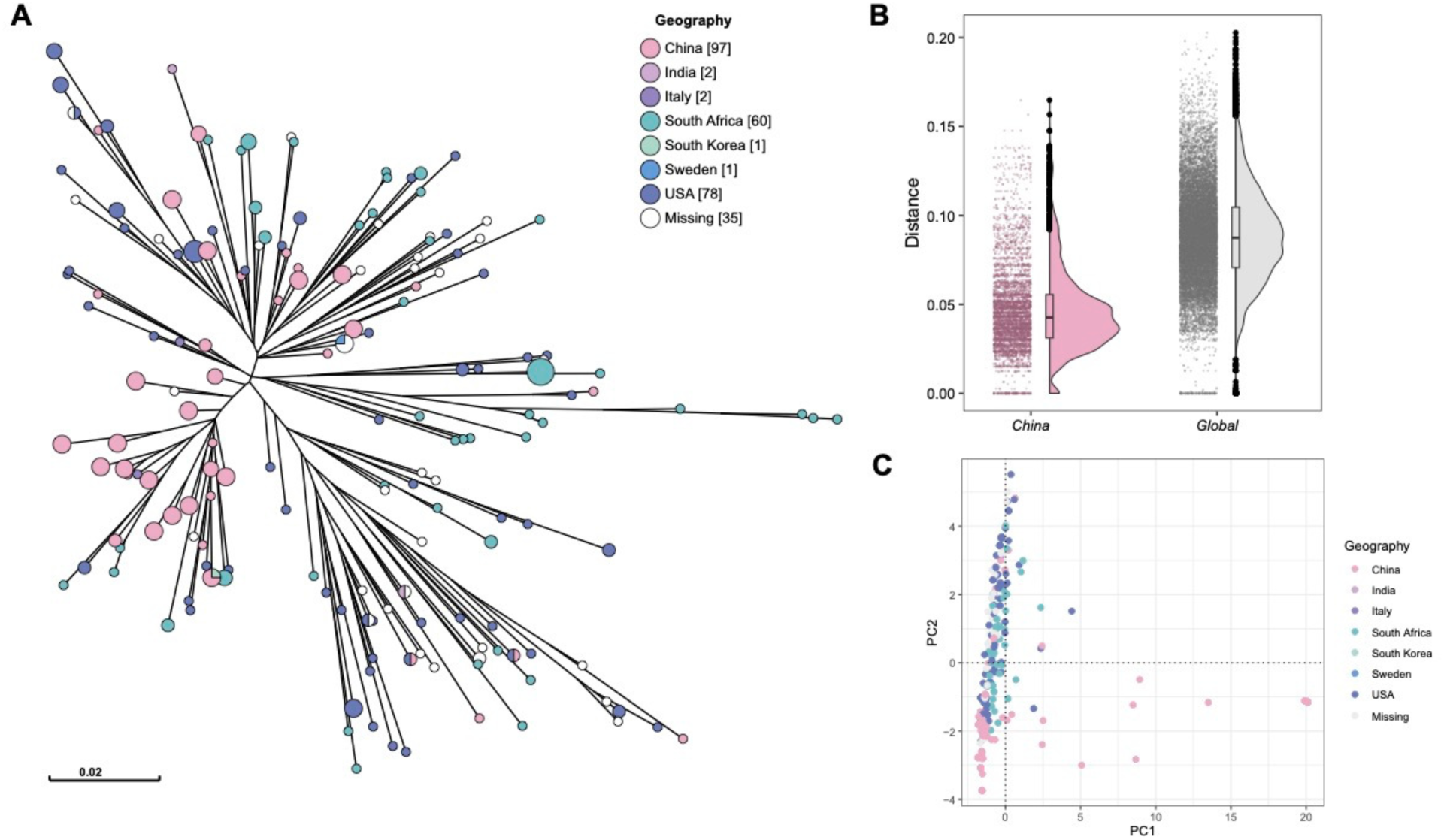
Analysis of the core genome of *Lactobacillus iners* A. Maximum likelihood phylogenetic tree of *Lactobacillus iners* (Includes *Lactobacillus iners* genomes obtained from this study and the NCBI public database). B. PCA analysis constructing a two-dimensional scatter plot of the core genome of *Lactobacillus iners* based on SNPs: C. Distribution of distances among *Lactobacillus iners* Strains (according to Figure A).

A similar trend was also observed in the pan-genome (Fig. 6). Chinese strains formed a loose cluster in the PCA plot and were separated into multiple distinct clusters in the gene content phylogeny (Fig. 6C). Separation of samples from North and South was still observed (p < 0.03), each intermingled with genomes from other countries.

**Figure 6.**
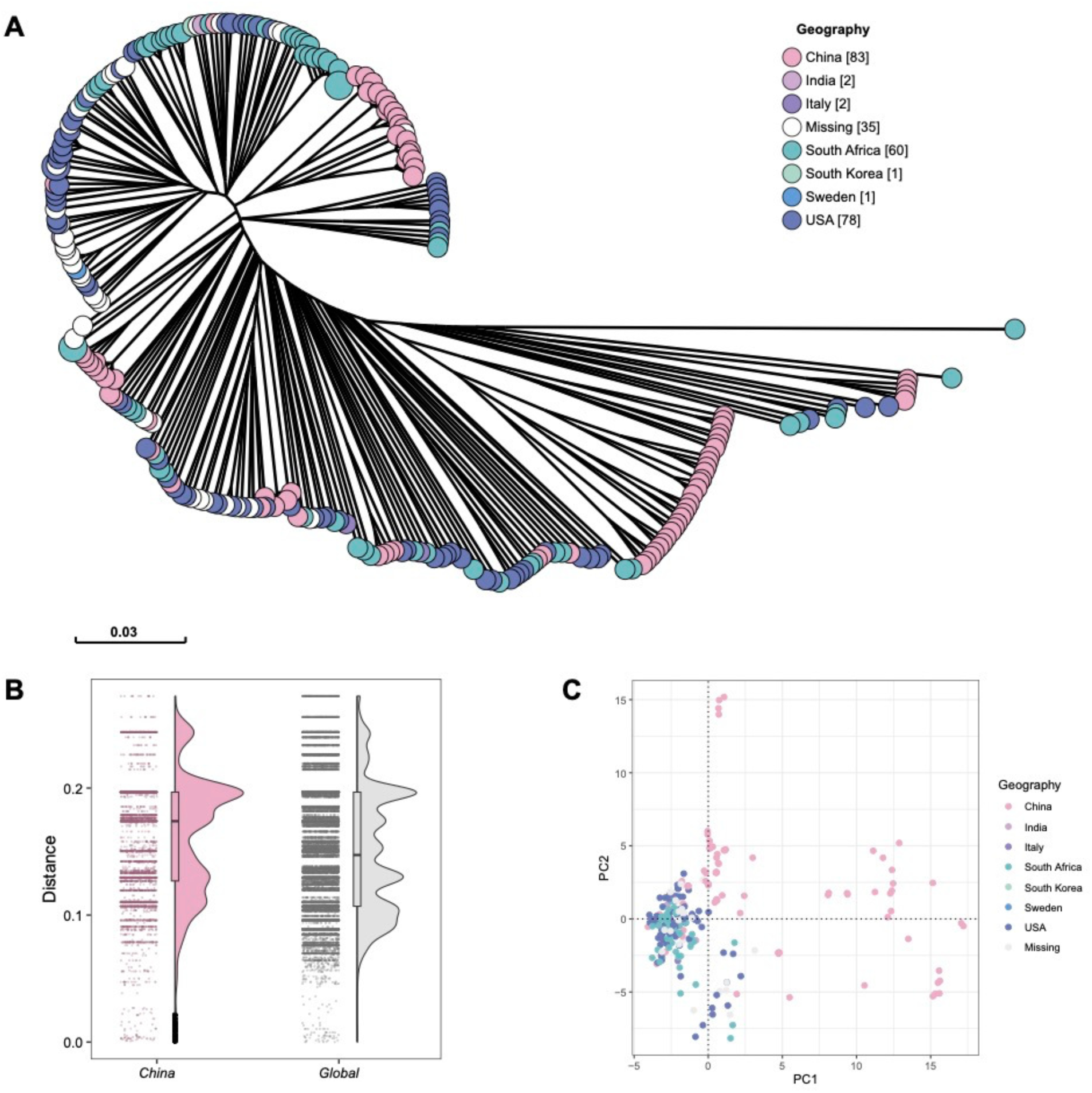
Pan-Genomic Analysis of *Lactobacillus crispatus* A Pan Genome tree of *Lactobacillus iners* (calculate the Bray-Curtis dissimilarity index between samples based on the pangenome gene presence/absence matrix and perform sample clustering analysis using the UPGMA method). B. Distribution of pangenome distances among Lactobacillus iners Strains(according to Figure A). C. PCA analysis constructing a two-dimensional scatter plot of the Lactobacillus iners pan genome based on the presence/absence of pan genome genes.

A Wilcoxon rank-sum test was conducted on the pan-genome of the *L. iners* strains to extract differential genes (p > 0.05), which were mostly related to protein transport and metabolism. Specifically, we observed a set of CRISPR system-related genes and the hsdR gene, which were both associated with phage resistence.

Finally, *G. vaginalis*, the predominant species in CST-VI-B, was genetically the most diverse of the three, and had been separated into at least four potential species (SGB) in MetaPhlan4 and GTDB (Fig. 7). The majority of *G. vaginalis* in China were from SGB17307 (Fig. 7 C). Unlike *Lactobacillus*, the Chinese samples did not form clear clusters within this species but were intermixed with international samples, indicating a complex evolutionary history of *G. vaginalis*.

**Figure 7.**
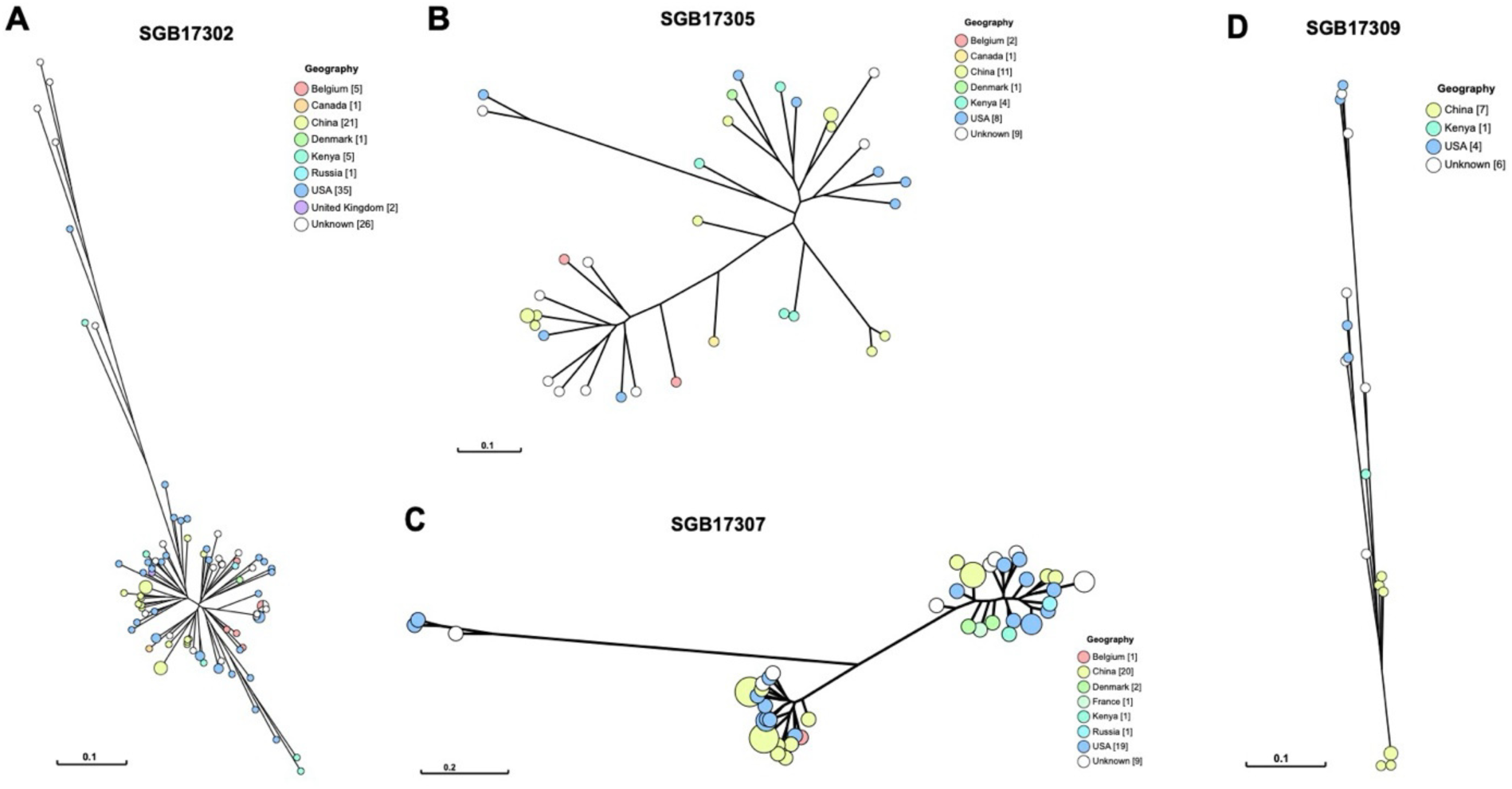
Phylogenetic Tree of Vaginal *Gardnerella* at the Strain Level Construct a phylogenetic tree of the strains using reconstructed markers: A. Reference genome reconstructed strain numbered SGB17302. B. Reference genome reconstructed strain numbered SGB17305. C. Reference genome reconstructed strain numbered SGB17307. D. Reference genome reconstructed strain numbered SGB17309.

## Discussion

This study offers novel insights into the geographic-specific evolution of the vaginal microbiota in Chinese women, with a particular focus on *Lactobacillus crispatus*, *Lactobacillus iners*, and *Gardnerella vaginalis*. The findings not only expand the existing knowledge on microbial diversity in different populations but also highlight the evolutionary dynamics shaped by both environmental and host factors (5, 22, 23).

Our analysis revealed geographic patterns in the distribution of community state types (CSTs) within the Chinese population. *L. crispatus*-dominated CST-I was more prevalent in northern regions, while *L. iners*-dominated CST-III showed higher prevalence in southern regions. These findings suggest that geographic location, along with diet and other lifestyle factors, plays a significant role in shaping the vaginal microbiota (20, 24–27).

Moreover, the study highlights the dynamic nature of the vaginal microbiota across different phases of the menstrual cycle. While most individuals maintained consistent CSTs, some shifted from *L. iners* to *L. crispatus* or from *G. vaginalis* to *L. iners*, suggesting that microbial communities can transition towards healthier states, influenced by hormonal and environmental factors (8, 28–33). Conversely, the bacterial vaginosis (BV) likely represented a dysfunctional state in the vagina microbiome, caused by the invasion of bacteria from other tracts (34–37).

Our results demonstrate that *L. crispatus* was a recent addition to the vaginal microbiota in China. The reduced genetic diversity and clustering of Chinese strains at the terminal branches of the phylogenetic tree, combined with their limited SNP variations, suggest a rapid and recent spread across the population. Phylogenetic analysis revealed that *L. crispatus* strains in China are most likely derived from a vaginal-associated population (VAP), distinguishing them from gut-associated populations (GAP) (38). This separation points to the adaptation of *L. crispatus* from its ancestral gut environment to the vaginal niche, reflecting recent colonization in human history (39). The divergence of the *L. crispatus* strains in China likely coincided with shifts in dietary habits, particularly the consumption of fermented dairy products, which may have facilitated its introduction into the vaginal microbiota from the gut (40).

This observation aligns with previous reports indicating a more recent colonization of *L. crispatus* in human history (39) compared to *L. iners* and *Gardnerella*, both of which are more deeply rooted in the vaginal microbiome of diverse populations. Our data suggest that *L. crispatus* may have entered northern China first, where dietary practices rich in fermented dairy products could have promoted its establishment and eventual southward spread.

In contrast to *L. crispatus*, *L. iners* displayed a higher level of genetic diversity, indicative of its longer residence within the vaginal microbiota of Chinese women. The evolutionary history of *L. iners* appears to date back further (19). The geographic specificity observed in the Chinese strains, with a notable division between northern and southern regions, points to region-specific selective pressures that have shaped the evolution of this species. The genetic diversity within Chinese *L. iners* strains, along with their distinct separation from international strains, underscores the influence of unique host environments and interactions within the vaginal microbiome (41).

The phylogenetic analysis of *Gardnerella vaginalis* in this study revealed significant genetic diversity, supporting the notion that this species has undergone complex evolutionary dynamics, potentially including horizontal gene transfer (42). Unlike *Lactobacillus*, the Chinese *G. vaginalis* strains did not form distinct clusters but were interspersed with international strains, suggesting a more interconnected and fluid evolutionary history (43). The intermingling of strains indicates that *G. vaginalis* has likely experienced frequent genetic exchanges across populations, which may contribute to its role in BV and other dysbiosis-related conditions (44).

The evolutionary insights gained from this study have important implications for understanding the role of the vaginal microbiota in women’s health. The rapid spread of *L. crispatus* across China suggests that this species may be beneficial in promoting vaginal health, given its association with low vaginal pH and protection against pathogens (6). In contrast, the high diversity and frequent horizontal gene transfers in *G. vaginalis* underline its potential to cause dysbiosis, particularly in regions where it is more prevalent (34).

Understanding the evolutionary dynamics of these key vaginal bacteria provides a foundation for developing targeted therapeutic interventions aimed at restoring and maintaining a healthy vaginal microbiome. The identification of region-specific microbiota patterns also raises the possibility of tailoring interventions based on geographic and population-specific microbiome characteristics.

## Conclusions

In conclusion, this study provides a comprehensive phylogenetic and genomic analysis of the vaginal microbiota in Chinese women, uncovering the recent introduction of *L. crispatus* and the long-standing presence of *L. iners* and *Gardnerella vaginalis*. The geographic-specific evolution of these species, coupled with their varied roles in vaginal health and disease, underscores the need for further research to explore the factors driving microbial community structure in different populations. These findings provide a pathway for personalized approaches to managing vaginal health based on regional microbiome characteristics.

## Methods

### Study population

In this study, from November 2022 to October 2023, 120 vaginal samples were collected nationwide from 30 women of reproductive age with regular menstrual cycles during four menstrual phases: follicular phase, pre-ovulation, post-ovulation, and luteal phase. The participants were from Beijing (n=5), Henan (n=7), Liaoning (n=7), Zhejiang (n=5), and Xinjiang (n=6).

Inclusion criteria include women aged 18 to 45 years with a history of sexual activity, not pregnant, and who have not taken any antibiotic or antimycotic compounds in the past 30 days, while exclusion criteria include age over 45 years, post-menopause, and antibiotics used within 1 month of enrolling in this study.

### Sample collection

After obtaining informed consent, the vaginal swab was obtained from the posterior fornix by physicians using MUNKCARE (Changzhou Munk Foam Technology Co.,Ltd). The first swabs were stored in 1 mL of ribonucleic acid (RNA) Storage Reagent (Invitrogen) at room temperature until transported to the laboratory, where they were stored at −80℃. And each participant completed a questionnaire on lifestyle, dietary habits, reproductive history, sexual lifestyle, and other information. This study was approved by the Ethics Committee of Peking Union Medical College Hospital (KT-2255)

### DNA Extraction

Swab samples were thawed on ice before analysis and vortexed vigorously for 5 min to resuspend cells. 1mL aliquots were transferred to sterile 1.5mL tubes and stored on ice. Cell lysis was initiated by adding 180μL of lysozyme to sample (10mg/mL, Vazyme). After 0.5h incubation at 37℃, cells were disrupted by mixing with bleached and rinsed 0.1mm-diameter zirconia beads for 3 min at room temperature, with 6m/s oscillations for 30 seconds followed by a 30 second rest period per cycle for a total of 6 cycles in a TGrinder H24R (TIANGEN Products). The resulting crude lysate was processed using FastPure Bacterial deoxynucleic acid (DNA) Isolation Mini Kit (Vazyme) according to the manufacturer’s instructions for processing of crude lysates. The samples were eluted with 40μL of ribonuclease (RNase)-free water into separate tubes. The DNA concentrations in the samples were measured using the Qubit® dsDNA HS Assay Kit from Molecular Probes (Invitrogen).

### Metagenome Library Construction and Sequencing

Sequencing libraries were prepared using the NadPrep® EZ DNA Library Preparation Module (Illumina®) following a modification of the manufacturer’s protocol: DNA was purified between enzymatic reactions and library size selection was performed with AMPure XP beads (Beckman Coulter Genomics). The library concentrations were measured using the Qubit® dsDNA HS Assay Kit from Molecular Probes (Invitrogen) and library quality was checked using an Agilent DNA 1000 Reagent from Agilent 4150 TapeStation. Library sequencing was performed using the Illumina Novaseq 6000 platform (150 bp paired end mode, 8 Gbp per sample).

### Species Classification

Species classification and relative abundance in the metagenome were obtained using MetaPhlAn4 with default parameters (45). MetaPhlAn4 integrates information from 1.01 million prokaryotic reference genomes and metagenome-assembled genomes, defining unique marker genes for 26,970 species-level genome bins to provide a more comprehensive metagenomic classification analysis.

### CST Type Assignment

Community State Types (CST) are clusters of community samples with similar microbial taxonomic compositions. Species abundance results were clustered using uniform manifold approximation and projection (UMAP), and CST types were defined for the samples based on the dominant species in the obtained clusters.

### Genomic and Pan-Genomic Analysis

Sequencing reads for each strain were quality trimmed using EtoKi prepare, and high-quality sequences were further assembled into contigs using SPAdes V3.13 via EtoKi assembler (46). Genes in each assembled genome were predicted and annotated using PROKKA 1.14.6 (47), with detailed functional predictions performed using eggnog-mapper v2 (48). Multiple sequence alignments for each species were generated using the EtoKi align module, and maximum likelihood (ML) phylogenetic trees were constructed using IQTree v1.6.12 implemented in EtoKi phylo(49).

Using Phylogeny Enhanced Pipeline for PAN-genome (PEPPAN), construct the presence/absence matrix of the pangenome genes for *Lactobacillus crispatus* and *Lactobacillus iners* reference genomes(50). Extract the core genome to serve as the reference sequence and estimate the coverage of strains in each sample. Retain samples with a coverage of greater than or equal to 4 and extract genes with coverage greater than the average coverage plus 0.2 for the presence/absence matrix of the pangenome.

Analyze the presence/absence matrix of the pangenome using the vegdist function from the vegan package in R to calculate the Bray-Curtis dissimilarity index between samples. Then, use the upgma function from the phangorn package to perform hierarchical clustering analysis of the samples. Visualize the results in grapetree(51).

### Population dynamics of *Lactobacillus crispatus*

We used rhierbaps with default parameters to divide the VAP into four clusters (52). To infer the population dynamics of the VAP-D cluster, we conducted analyses using BEAST2 (53). A total of 4 alternative models were compared using nested sampling tests, and the model with optimal relaxed substitution rates and Bayesian skyline coalescents was selected based on the greatest Bayes factor. The evolutionary tree was visualized using iTOL (54).

### Construction of the Phylogenetic Tree of Vaginal Gardnerella

Phylogenetic trees were generated using StrainPhlAn4 (v4.0.6) (55). This method utilizes unique clade-specific marker genes used by MetaPhlAn4 to reconstruct the sample-specific gene composition of individual strains. Reconstructed markers with <50% coverage were discarded. Consensus sequences were trimmed by removing the first and last 50 bases due to lower coverage at terminal positions caused by read mapping limitations to truncated sequences. Next, markers appearing in <50% of samples were deleted. Samples with <50% marker percentages were removed from the alignment.Phylogenetic trees were produced by StrainPhlAn4 based on a comparison of marker genes.

## List of Abbreviations

CST: community state type
AI: artificial intelligence
RNA: ribonucleic acid
DNA: deoxynucleic acid
RNase: ribonuclease
UMAP: uniform manifold approximation and projection
ML: maximum likelihood
PEPPAN: Phylogeny Enhanced Pipeline for PAN-genome
SDI: Simpson diversity indices
MAG: metagenome-assembled genome
PCoA: principal coordinates analysis
EtoKi: Enterobase ToolKit
SNP: single nucleotide polymorphism
VAP: vagina-associted populations
GAP: gut-associated populations
PCA: principle component analysis
SGB: Species Genome Bin
GTDB: Genome Taxonomy Database
BV: bacterial vaginosis.

## Declarations

Ethics approval and consent to participate

This study was approved by the Ethics Committee of Peking Union Medical College Hospital (approval No. ZS-2255) and informed consent was obtained from all participants.

## Consent for publication

Not applicable.

## Availability of data and material

The raw metagenomic sequencing data generated during this study have been uploaded to the National Genomics Data Center (NGDC) under accession number PRJCA031189. These datasets are publicly available through the NGDC GSA database and can be accessed at https://ngdc.cncb.ac.cn/gsub.

## Competing interests

The authors report there are no competing interests to declare.

## Funding

This study was supported by the National High Level Hospital Clinical Research Funding (2022-PUMCH-A-114), and Beijing Natural Science Foundation (L232074).

## Authors’ contributions

HY and LZ conceived the project. HY, LJH, JTH, CYZ and LLL collected the samples and information. YNL performe statistical data analysis. ZMZ introduced enrichment scores in the analysis and helped interpret the data. All authors read, revised, and approved the final manuscript.

## Acknowledgements

We thank the platforms of National Clinical Research Center for Obstetric & Gynecologic Diseases, Peking Union Medical College Hospital and Key Laboratory of Alkene-carbon Fibres-based Technology & Application for Detection of Major Infectious Diseases, Suzhou Medical College, Soochow University for technical help.

## References

1. Ma B, Forney LJ, Ravel J. 2012. Vaginal microbiome: rethinking health and disease. Annu Rev Microbiol 66:371–89.

2. Greenbaum S, Greenbaum G, Moran-Gilad J, Weintraub AY. 2019. Ecological dynamics of the vaginal microbiome in relation to health and disease. Am J Obstet Gynecol 220:324–335.

3. Anonymous. 2017. Comprehensive and Integrated Genomic Characterization of Adult Soft Tissue Sarcomas. Cell 171:950–965.e28.

4. Balle C, Konstantinus IN, Jaumdally SZ, Havyarimana E, Lennard K, Esra R, Barnabas SL, Happel AU, Moodie Z, Gill K, Pidwell T, Karaoz U, Brodie E, Maseko V, Gamieldien H, Bosinger SE, Myer L, Bekker LG, Passmore JS, Jaspan HB. 2020. Hormonal contraception alters vaginal microbiota and cytokines in South African adolescents in a randomized trial. Nat Commun 11:5578.

5. Noyes N, Cho KC, Ravel J, Forney LJ, Abdo Z. 2018. Associations between sexual habits, menstrual hygiene practices, demographics and the vaginal microbiome as revealed by Bayesian network analysis. PLoS One 13:e0191625.

6. Ravel J, Gajer P, Abdo Z, Schneider GM, Koenig SS, McCulle SL, Karlebach S, Gorle R, Russell J, Tacket CO, Brotman RM, Davis CC, Ault K, Peralta L, Forney LJ. 2011. Vaginal microbiome of reproductive-age women. Proc Natl Acad Sci U S A 108 Suppl 1:4680–7.

7. Kaur H, Merchant M, Haque MM, Mande SS. 2020. Crosstalk Between Female Gonadal Hormones and Vaginal Microbiota Across Various Phases of Women’s Gynecological Lifecycle. Front Microbiol 11:551.

8. Song SD, Acharya KD, Zhu JE, Deveney CM, Walther-Antonio MRS, Tetel MJ, Chia N. 2020. Daily Vaginal Microbiota Fluctuations Associated with Natural Hormonal Cycle, Contraceptives, Diet, and Exercise. mSphere 5.

9. France MT, Ma B, Gajer P, Brown S, Humphrys MS, Holm JB, Waetjen LE, Brotman RM, Ravel J. 2020. VALENCIA: a nearest centroid classification method for vaginal microbial communities based on composition. Microbiome 8:166.

10. Li T, Liu Z, Zhang X, Chen X, Wang S. 2019. Local Probiotic Lactobacillus crispatus and Lactobacillus delbrueckii Exhibit Strong Antifungal Effects Against Vulvovaginal Candidiasis in a Rat Model. Front Microbiol 10:1033.

11. Ojala T, Kankainen M, Castro J, Cerca N, Edelman S, Westerlund-Wikström B, Paulin L, Holm L, Auvinen P. 2014. Comparative genomics of Lactobacillus crispatus suggests novel mechanisms for the competitive exclusion of Gardnerella vaginalis. BMC Genomics 15:1070.

12. Chee WJY, Chew SY, Than LTL. 2020. Vaginal microbiota and the potential of Lactobacillus derivatives in maintaining vaginal health. Microb Cell Fact 19:203.

13. Muzny CA, Łaniewski P, Schwebke JR, Herbst-Kralovetz MM. 2020. Host-vaginal microbiota interactions in the pathogenesis of bacterial vaginosis. Curr Opin Infect Dis 33:59–65.

14. Petrova MI, Reid G, Vaneechoutte M, Lebeer S. 2017. Lactobacillus iners: Friend or Foe? Trends Microbiol 25:182–191.

15. Gupta S, Kakkar V, Bhushan I. 2019. Crosstalk between Vaginal Microbiome and Female Health: A review. Microb Pathog 136:103696.

16. Zhou X, Brown CJ, Abdo Z, Davis CC, Hansmann MA, Joyce P, Foster JA, Forney LJ. 2007. Differences in the composition of vaginal microbial communities found in healthy Caucasian and black women. Isme j 1:121–33.

17. Roachford OSE, Alleyne AT, Nelson KE. 2022. Insights into the vaginal microbiome in a diverse group of women of African, Asian and European ancestries. PeerJ 10:e14449.

18. Fettweis JM, Brooks JP, Serrano MG, Sheth NU, Girerd PH, Edwards DJ, Strauss JF, The Vaginal Microbiome C, Jefferson KK, Buck GA. 2014. Differences in vaginal microbiome in African American women versus women of European ancestry. Microbiology (Reading) 160:2272–2282.

19. Nielsen R, Akey JM, Jakobsson M, Pritchard JK, Tishkoff S, Willerslev E. 2017. Tracing the peopling of the world through genomics. Nature 541:302–310.

20. Morsli M, Gimenez E, Magnan C, Salipante F, Huberlant S, Letouzey V, Lavigne JP. 2024. The association between lifestyle factors and the composition of the vaginal microbiota: a review. Eur J Clin Microbiol Infect Dis 43:1869–1881.

21. Zhou Z, Alikhan NF, Mohamed K, Fan Y, Achtman M. 2020. The EnteroBase user’s guide, with case studies on Salmonella transmissions, Yersinia pestis phylogeny, and Escherichia core genomic diversity. Genome Res 30:138–152.

22. Si J, You HJ, Yu J, Sung J, Ko G. 2017. Prevotella as a Hub for Vaginal Microbiota under the Influence of Host Genetics and Their Association with Obesity. Cell Host Microbe 21:97–105.

23. Gliniewicz K, Schneider GM, Ridenhour BJ, Williams CJ, Song Y, Farage MA, Miller K, Forney LJ. 2019. Comparison of the Vaginal Microbiomes of Premenopausal and Postmenopausal Women. Front Microbiol 10:193.

24. Gupta VK, Paul S, Dutta C. 2017. Geography, Ethnicity or Subsistence-Specific Variations in Human Microbiome Composition and Diversity. Front Microbiol 8:1162.

25. Marconi C, El-Zein M, Ravel J, Ma B, Lima MD, Carvalho NS, Alves RRF, Parada C, Leite SHM, Giraldo PC, Gonçalves AK, Franco EL, Silva MG. 2020. Characterization of the Vaginal Microbiome in Women of Reproductive Age From 5 Regions in Brazil. Sex Transm Dis 47:562–569.

26. Rosen EM, Martin CL, Siega-Riz AM, Dole N, Basta PV, Serrano M, Fettweis J, Wu M, Sun S, Thorp JM, Jr., Buck G, Fodor AA, Engel SM. 2022. Is prenatal diet associated with the composition of the vaginal microbiome? Paediatr Perinat Epidemiol 36:243–253.

27. Miller EA, Beasley DE, Dunn RR, Archie EA. 2016. Lactobacilli Dominance and Vaginal pH: Why Is the Human Vaginal Microbiome Unique? Front Microbiol 7:1936.

28. Gajer P, Brotman RM, Bai G, Sakamoto J, Schütte UM, Zhong X, Koenig SS, Fu L, Ma ZS, Zhou X, Abdo Z, Forney LJ, Ravel J. 2012. Temporal dynamics of the human vaginal microbiota. Sci Transl Med 4:132ra52.

29. Uchihashi M, Bergin IL, Bassis CM, Hashway SA, Chai D, Bell JD. 2015. Influence of age, reproductive cycling status, and menstruation on the vaginal microbiome in baboons (Papio anubis). Am J Primatol 77:563–78.

30. Krog MC, Hugerth LW, Fransson E, Bashir Z, Nyboe Andersen A, Edfeldt G, Engstrand L, Schuppe-Koistinen I, Nielsen HS. 2022. The healthy female microbiome across body sites: effect of hormonal contraceptives and the menstrual cycle. Hum Reprod 37:1525–1543.

31. Xu J, Bian G, Zheng M, Lu G, Chan WY, Li W, Yang K, Chen ZJ, Du Y. 2020. Fertility factors affect the vaginal microbiome in women of reproductive age. Am J Reprod Immunol 83:e13220.

32. Wessels JM, Felker AM, Dupont HA, Kaushic C. 2018. The relationship between sex hormones, the vaginal microbiome and immunity in HIV-1 susceptibility in women. Dis Model Mech 11.

33. Farage MA, Miller KW, Sobel JD. 2010. Dynamics of the Vaginal Ecosystem—Hormonal Influences. Infectious Diseases: Research and Treatment 3:IDRT.S3903.

34. Kaambo E, Africa C, Chambuso R, Passmore JS. 2018. Vaginal Microbiomes Associated With Aerobic Vaginitis and Bacterial Vaginosis. Front Public Health 6:78.

35. Chen X, Lu Y, Chen T, Li R. 2021. The Female Vaginal Microbiome in Health and Bacterial Vaginosis. Front Cell Infect Microbiol 11:631972.

36. Ling Z, Kong J, Liu F, Zhu H, Chen X, Wang Y, Li L, Nelson KE, Xia Y, Xiang C. 2010. Molecular analysis of the diversity of vaginal microbiota associated with bacterial vaginosis. BMC Genomics 11:488.

37. Africa CW, Nel J, Stemmet M. 2014. Anaerobes and bacterial vaginosis in pregnancy: virulence factors contributing to vaginal colonisation. Int J Environ Res Public Health 11:6979–7000.

38. Zhang Q, Zhang L, Ross P, Zhao J, Zhang H, Chen W. 2020. Comparative Genomics of Lactobacillus crispatus from the Gut and Vagina Reveals Genetic Diversity and Lifestyle Adaptation. Genes (Basel) 11.

39. Tortelli BA, Lewis AL, Fay JC. 2021. The structure and diversity of strain-level variation in vaginal bacteria. Microb Genom 7.

40. G BM, M GS, Marconi C. 2023. Milk and Dairy Consumption and Its Relationship With Abundance of Lactobacillus crispatus in the Vaginal Microbiota: Milk Intake and Vaginal Lactobacillus. J Low Genit Tract Dis 27:280–285.

41. Pavlova SI, Kilic AO, Kilic SS, So JS, Nader-Macias ME, Simoes JA, Tao L. 2002. Genetic diversity of vaginal lactobacilli from women in different countries based on 16S rRNA gene sequences. J Appl Microbiol 92:451–9.

42. France MT, Mendes-Soares H, Forney LJ. 2016. Genomic Comparisons of Lactobacillus crispatus and Lactobacillus iners Reveal Potential Ecological Drivers of Community Composition in the Vagina. Appl Environ Microbiol 82:7063–7073.

43. Tarracchini C, Lugli GA, Mancabelli L, Milani C, Turroni F, Ventura M. 2020. Assessing the Genomic Variability of Gardnerella vaginalis through Comparative Genomic Analyses: Evolutionary and Ecological Implications. Appl Environ Microbiol 87.

44. Ahmed A, Earl J, Retchless A, Hillier SL, Rabe LK, Cherpes TL, Powell E, Janto B, Eutsey R, Hiller NL, Boissy R, Dahlgren ME, Hall BG, Costerton JW, Post JC, Hu FZ, Ehrlich GD. 2012. Comparative genomic analyses of 17 clinical isolates of Gardnerella vaginalis provide evidence of multiple genetically isolated clades consistent with subspeciation into genovars. J Bacteriol 194:3922–37.

45. Blanco-Míguez A, Beghini F, Cumbo F, McIver LJ, Thompson KN, Zolfo M, Manghi P, Dubois L, Huang KD, Thomas AM, Nickols WA, Piccinno G, Piperni E, Punčochář M, Valles-Colomer M, Tett A, Giordano F, Davies R, Wolf J, Berry SE, Spector TD, Franzosa EA, Pasolli E, Asnicar F, Huttenhower C, Segata N. 2023. Extending and improving metagenomic taxonomic profiling with uncharacterized species using MetaPhlAn 4. Nat Biotechnol 41:1633–1644.

46. Bankevich A, Nurk S, Antipov D, Gurevich AA, Dvorkin M, Kulikov AS, Lesin VM, Nikolenko SI, Pham S, Prjibelski AD, Pyshkin AV, Sirotkin AV, Vyahhi N, Tesler G, Alekseyev MA, Pevzner PA. 2012. SPAdes: a new genome assembly algorithm and its applications to single-cell sequencing. J Comput Biol 19:455–77.

47. Seemann T. 2014. Prokka: rapid prokaryotic genome annotation. Bioinformatics 30:2068–9.

48. Cantalapiedra CP, Hernández-Plaza A, Letunic I, Bork P, Huerta-Cepas J. 2021. eggNOG-mapper v2: Functional Annotation, Orthology Assignments, and Domain Prediction at the Metagenomic Scale. Mol Biol Evol 38:5825–5829.

49. Nguyen LT, Schmidt HA, von Haeseler A, Minh BQ. 2015. IQ-TREE: a fast and effective stochastic algorithm for estimating maximum-likelihood phylogenies. Mol Biol Evol 32:268–74.

50. Zhou Z, Charlesworth J, Achtman M. 2020. Accurate reconstruction of bacterial pan- and core-genomes with PEPPAN. bioRxiv doi:10.1101/2020.01.03.894154:2020.01.03.894154.

51. Zhou Z, Alikhan NF, Sergeant MJ, Luhmann N, Vaz C, Francisco AP, Carriço JA, Achtman M. 2018. GrapeTree: visualization of core genomic relationships among 100,000 bacterial pathogens. Genome Res 28:1395–1404.

52. Tonkin-Hill G, Lees JA, Bentley SD, Frost SDW, Corander J. 2018. RhierBAPS: An R implementation of the population clustering algorithm hierBAPS. Wellcome Open Res 3:93.

53. Bouckaert R, Heled J, Kühnert D, Vaughan T, Wu CH, Xie D, Suchard MA, Rambaut A, Drummond AJ. 2014. BEAST 2: a software platform for Bayesian evolutionary analysis. PLoS Comput Biol 10:e1003537.

54. Letunic I, Bork P. 2019. Interactive Tree Of Life (iTOL) v4: recent updates and new developments. Nucleic Acids Res 47:W256–w259.

55. Truong DT, Tett A, Pasolli E, Huttenhower C, Segata N. 2017. Microbial strain-level population structure and genetic diversity from metagenomes. Genome Res 27:626–638.

